# FAM83H couples keratin organization to Notch signaling during epidermal morphogenesis

**DOI:** 10.64898/2026.05.26.727959

**Authors:** Mageshi Kamaraj, Justin K. Amakor, Betul Melike Ogan, Grace Qian, Kyle A. Jacobs, Arvind Arul Nambi Rajan, Radislav Sedlacek, Torsten Wittmann, Erica J. Hutchins, Jana Balounová, Matthew L. Kutys

## Abstract

Keratin intermediate filaments are essential for epidermal integrity, yet how keratin organization is coupled to the cell fate signaling that coordinates keratinocyte differentiation during epidermal morphogenesis remains poorly understood. Here, we identify FAM83H as a previously unrecognized regulator that links keratin cytoskeletal organization to epidermal cell fate decisions. Using mouse models and a human 3D microphysiological epidermis model, we show that loss of FAM83H disrupts epidermal architecture by impairing basal keratinocyte differentiation, organization, and cell-cell adhesion. Single-cell transcriptomic analysis of 3D epidermal tissues identified Notch signaling as prominently associated with cell populations lost upon FAM83H depletion. Mechanistically, FAM83H localizes to cell-cell junctions, where it organizes keratin filaments and desmosome integrity. Loss of FAM83H disrupts a desmoplakin-keratin-Notch1 complex at junctions, impairing Notch1 proteolytic activation and proper keratinocyte differentiation. Together, our work identifies FAM83H as a key regulator of epidermal morphogenesis that couples keratin cytoskeletal architecture to Notch1 signaling, and positions keratin-associated proteins as active participants in the epithelial fate decisions that govern epidermal homeostasis.

## Introduction

The epidermis is a stratified epithelium maintained by continuous self-renewal throughout adult life. Basal keratinocytes proliferate and subsequently stratify and differentiate to replenish suprabasal layers (Leary et al. 1992; Alberts et al. 2002; Fuchs and Raghavan 2002). Cells delaminate from the basal layer and migrate into the upper epidermis, enabled by loss of basement membrane contact and a cuboidal-to-squamous transition during which the cells undergo terminal differentiation (Liebig et al. 2009; Damen et al. 2021). This process is coordinated in part by biomechanical signaling at cell-cell and cell-extracellular matrix contacts that links cell fate to tissue architecture (Rübsam et al. 2017; Miroshnikova et al. 2018; Sahu et al. 2025; Villeneuve et al. 2026), and alteration of this coordination underlies several diseases including autoimmune conditions, psoriasis, and cancer (Iizuka et al. 2004; Segre 2006; Fuchs 2007). Still, the mechanisms that couple keratinocyte differentiation with mechanics and spatial organization remain incompletely understood.

During epidermal keratinocyte differentiation, expression of different keratin intermediate filament isoforms marks the distinct layers of the epidermis; basal cells characteristically express keratin (KRT) KRT5/KRT14 and suprabasal cells express KRT1/KRT10 (Blanpain and Fuchs 2009). Effective epidermal barrier function depends on keratin intermediate filaments (Kumar et al. 2015), but how keratins are linked to fate signaling associated with epidermal morphogenesis is not clear. In the epidermis, keratin mutations cause aberrant epidermal differentiation and lead to skin blistering diseases such as epidermolysis bullosa simplex, epidermolytic hyperkeratosis and epidermolytic palmoplantar keratoderma (Coulombe et al. 1991, 2009). This suggests that their functions may not be solely limited to mechanical support. Beyond structural roles, keratins have emerged as regulators of cell proliferation, differentiation and signaling (Alam et al. 2011; Nanes et al. 2024; Redmond et al. 2026), including modulation of Notch1 signaling and fate decisions in thymic and colonic epithelia (Santos et al. 2005; Lähdeniemi et al. 2017). However, how keratin organization might contribute to the coordination of keratinocyte fate and spatial organization is not known.

Family of Sequence Similarity 83H (FAM83H/SACK1H) is an understudied regulator of keratin filament organization initially identified through familial mutations causing amelogenesis imperfecta (Kim et al. 2008; Kuga et al. 2013; S. K. Wang et al. 2021). Broadly expressed across epithelial tissues including the epidermis (Karlsson et al. 2021; Ogan et al. 2026; “FAM83H Protein Expression Summary - The Human Protein Atlas,” n.d.), FAM83H expression is dysregulated in multiple cancers, with reduced levels in cutaneous squamous and basal cell carcinomas relative to normal epidermis (Tokuchi et al. 2021). *Fam83h* knockout mice exhibit epidermal abnormalities, including a scruffy coat and skin blistering, together implicating FAM83H in epidermal development and homeostasis (Shih-Kai Wang et al. 2015; Zheng et al. 2023; Ogan et al. 2026).

Like other FAM83 family members, FAM83H possesses a conserved N-terminal DUF1669 domain that mediates interactions with Casein Kinase 1 (CK1) isoforms to regulate their subcellular targeting (Fulcher et al. 2018). Distinct FAM83 family members scaffold CK1 to specific compartments to regulate processes such as spindle positioning and Wnt signaling (Fulcher et al. 2019; Dunbar et al. 2021; Wu et al. 2019). FAM83H is unique among family members in its association with keratin filaments via a C-terminal domain (Tachie-Menson et al. 2020), and both its loss and overexpression disrupt keratin organization across multiple epithelial cell types (Kuga et al. 2013, 2016; Tokuchi et al. 2021). Mechanistically, FAM83H has been proposed to scaffold CK1 to keratins to promote keratin phosphorylation and filament stabilization (Kuga et al. 2022), though, additional FAM83H domains and interactions remain largely unexplored.

In this study, we identify FAM83H as a critical regulator of epidermal morphogenesis in mice and in a biomimetic human 3D epidermal microphysiological model. FAM83H loss disrupts epidermal architecture via impaired basal cell differentiation. Single-cell RNA sequencing combined with biochemical analyses in keratinocytes reveals that FAM83H regulates epidermal morphogenesis through effects on keratin organization, desmosomes, and Notch1 signaling. Using domain-specific truncations, we identify that FAM83H co-localizes with desmoplakin (DSP) at desmosomes. We find that FAM83H is critical for the association of KRT5, DSP, and Notch1, and disruption of this complex via *FAM83H* deletion impairs Notch1 proteolytic activation during keratinocyte differentiation. Altogether, we demonstrate that FAM83H links keratin organization to Notch1 activation which is necessary for basal keratinocyte differentiation during epidermal morphogenesis.

## Results

### *Fam83h* deletion results in epidermal hyperplasia and impaired keratinocyte differentiation and cellular architecture

To investigate functions for *Fam83h* in epidermal morphogenesis, we utilized the CRISPR/Cas9-based approach to generate a *Fam83h* knockout mice within the IMPC effort (*Fam83h^em2^*^(*IMPC)Ccpcz*^, hereafter *Fam83h*^-/-^). As previously reported, *Fam83h^-/-^* mice have significant postnatal lethality and few mice survive for up to two-four weeks and exhibit abnormalities such as smaller size, hypoactivity, and skin defects such as sparse and scruffy coat, scaly skin, and blistering (Supplementary Fig. 1A) (Ogan et al. 2026; Shih-Kai Wang et al. 2015). Immunofluorescence staining revealed FAM83H is expressed in all layers of the epidermis (Supplementary Fig. 1B) and histopathological analysis of *Fam83h^-/-^* skin revealed epidermal hyperplasia and keratosis (Supplementary Fig. 1C).

To understand how loss of FAM83H causes epidermal hyperplasia, we performed immunofluorescence staining on sections of mouse back skin for basal (KRT14) and suprabasal (KRT10) keratins. In contrast to clearly delineated basal and suprabasal layers in wildtype littermates, *Fam83h^-/-^* skin was characterized by expanded KRT14 basal keratinocytes and an increase in dual positive KRT14 and KRT10 keratinocytes (Fig. 1A-C). Keratinocytes in *Fam83h^-/-^* epidermis multilayer but retain basal characteristics, including KRT14 expression and cuboidal cell morphology. Indeed, staining for filamentous actin (F-actin) revealed pronounced dysregulation of cell architecture across all epidermal layers in *Fam83h^-/-^*. Control epidermis exhibited basal cells transitioning to well-organized, thin squamous layers, whereas *Fam83h^-/-^*tissue lacked clearly stratified layers accompanied by cells lacking squamous morphology (Fig. 1A, D).

**Fig. 1.**
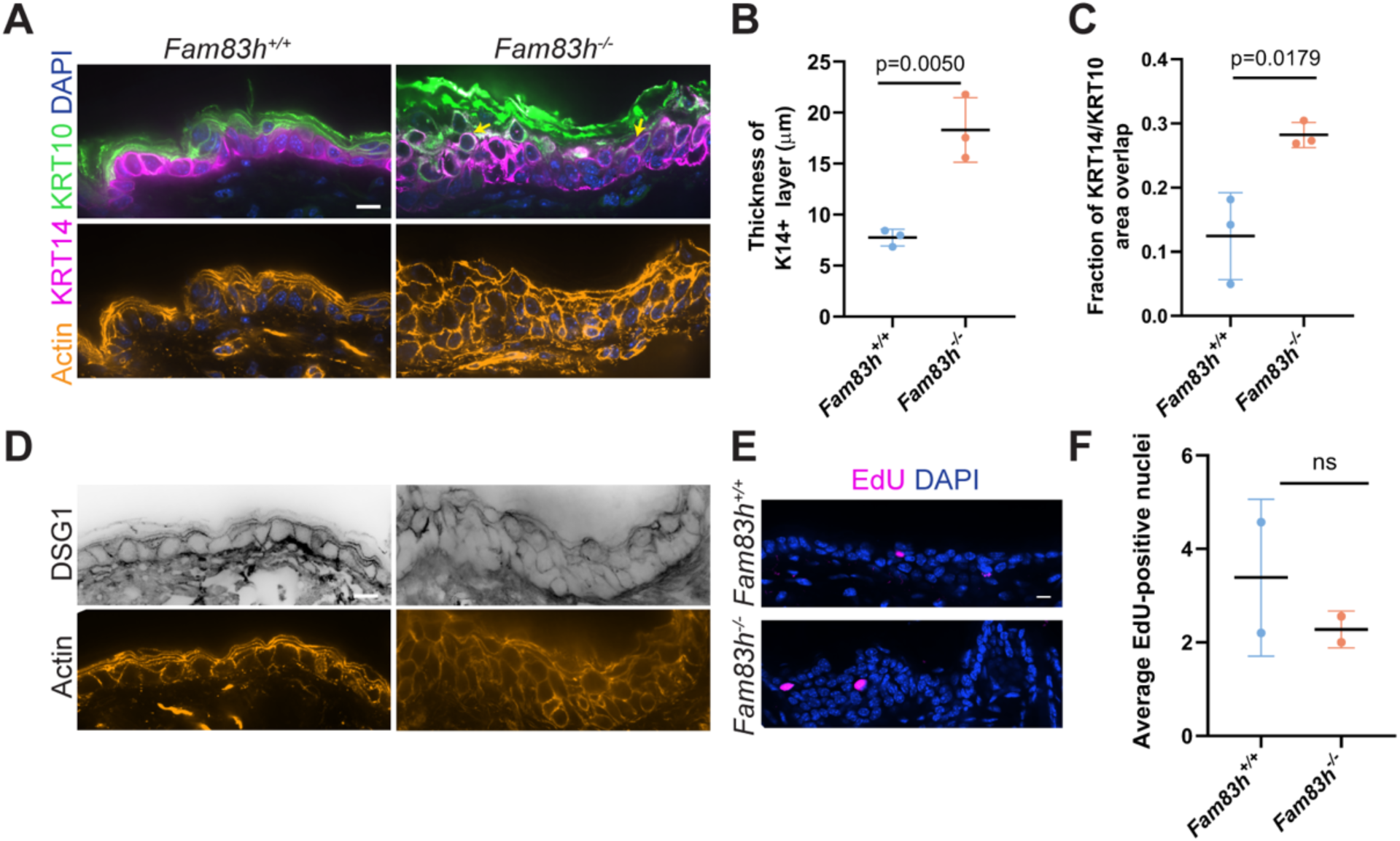
*Fam83h* deletion results in epidermal hyperplasia and impaired keratinocyte differentiation and cellular architecture. A) Representative z-slice micrographs of back skin sections from *Fam83h^+/+^* and *Fam83h^-/-^* mice at P16, immunostained for KRT14 (magenta) and KRT10 (green), phalloidin stained for F-actin (orange) and DAPI stained for nuclei (blue). Yellow arrows indicate a few KRT14+/KRT10+ cells. Scale bar, 10 μm. B) Quantification of basal layer thickness measured from the KRT14 channel. Measurements were taken across multiple images and averaged per mouse. N = 3 mice per group. C) Quantification of KRT14/KRT10 overlap, expressed as the fraction of overlapping area relative to total epidermal area. Measurements were taken across multiple images and averaged per mouse. N = 3 mice per group. D) Representative z-slice micrographs of back skin sections from *Fam83h^+/+^* and *Fam83h^-/-^*mice at P16, immunostained for DSG1 and phalloidin stained for F-actin (orange). Scale bar, 10 μm. E) Representative z-slice micrographs of back skin sections from *Fam83h^+/+^*and *Fam83h^-/-^* mice at P16 following EdU incorporation assay. EdU-positive nuclei (magenta) and DAPI stained for nuclei (blue). F) Quantification of EdU-positive nuclei per field of view. Measurements were taken across multiple images and averaged per mouse. N = 2 mice per group. In all plots, data represent mean ± SD. Statistical significance was determined by unpaired two-tailed student’s t-test. Exact p values are displayed on each plot; ns, non-significant.

Given the pronounced differences in keratinocyte morphology upon loss of FAM83H, we next investigated changes in cell-cell adherens and desmosome junctions. E-cadherin localization and intensity was not detectably different between *Fam83h*^+/+^ and *Fam83h^-/-^* epidermis (Supplementary Fig. 1D). Immunofluorescence staining of desmoglein-1 (DSG1) frequently revealed decreased junctional localization in *Fam83h^-/-^* epidermis relative to control (Fig. 1D), consistent with previous reports of desmosome defects in human ameloblastoma cells upon loss of Fam83h (Kuga et al. 2016). To assess the role of proliferation in *Fam83h^-/-^* basal cell expansion, *Fam83h*^+/+^ and *Fam83h*^⁻/⁻^ mice were injected with EdU prior to euthanasia and EdU incorporation analysis revealed no significant difference in EdU-positive nuclei (Fig. 1E, F). Given the broad epithelial defects in *Fam83h^-/-^* mice and the genetic connection to amelogenesis imperfecta, we examined the organization of another stratified tissue, the oral mucosa. Immunofluorescence staining of *Fam83h^-/-^* mucosal tissue similarly revealed basal keratinocyte expansion, aberrant cell size, and cells with dual KRT14 and KRT10 expression (Supplementary Fig. 1E). Thus, loss of *Fam83h* leads to defective epidermal morphogenesis through tissue hyperplasia that is driven by impaired basal keratinocyte differentiation and squamous cell morphology.

### Loss of FAM83H dysregulates basal keratinocyte differentiation and epidermal morphogenesis in a 3D human microphysiological system

To directly examine whether FAM83H plays a conserved role in human epidermal morphogenesis, we employed a 3D epidermal microphysiological system. This system allowed us to isolate keratinocyte-intrinsic effects independent of immune contributions. We utilized our previously characterized StrataChip microfluidic platform, which reproducibly captures keratinocyte fate and mechanical dynamics during epidermal morphogenesis in human skin equivalents generated from primary human dermal fibroblasts and epidermal keratinocytes (Ker-CT) (Amakor et al. 2026) (Fig. 2A). To assess the role of FAM83H, we generated keratinocytes constitutively expressing a FAM83H targeting shRNA hairpin (*FAM83H^KD^*), along with a nonspecific targeting control (NS). Keratinocytes were cultured on dermal equivalent hydrogels prior to introduction of an air-liquid interface, which induced stratification and differentiation of all four primary epidermal layers within 7 days. *In situ* immunostaining confirmed the efficient knockdown of FAM83H protein within 3D epidermal tissues (Supplementary Fig. 2A).

**Fig. 2.**
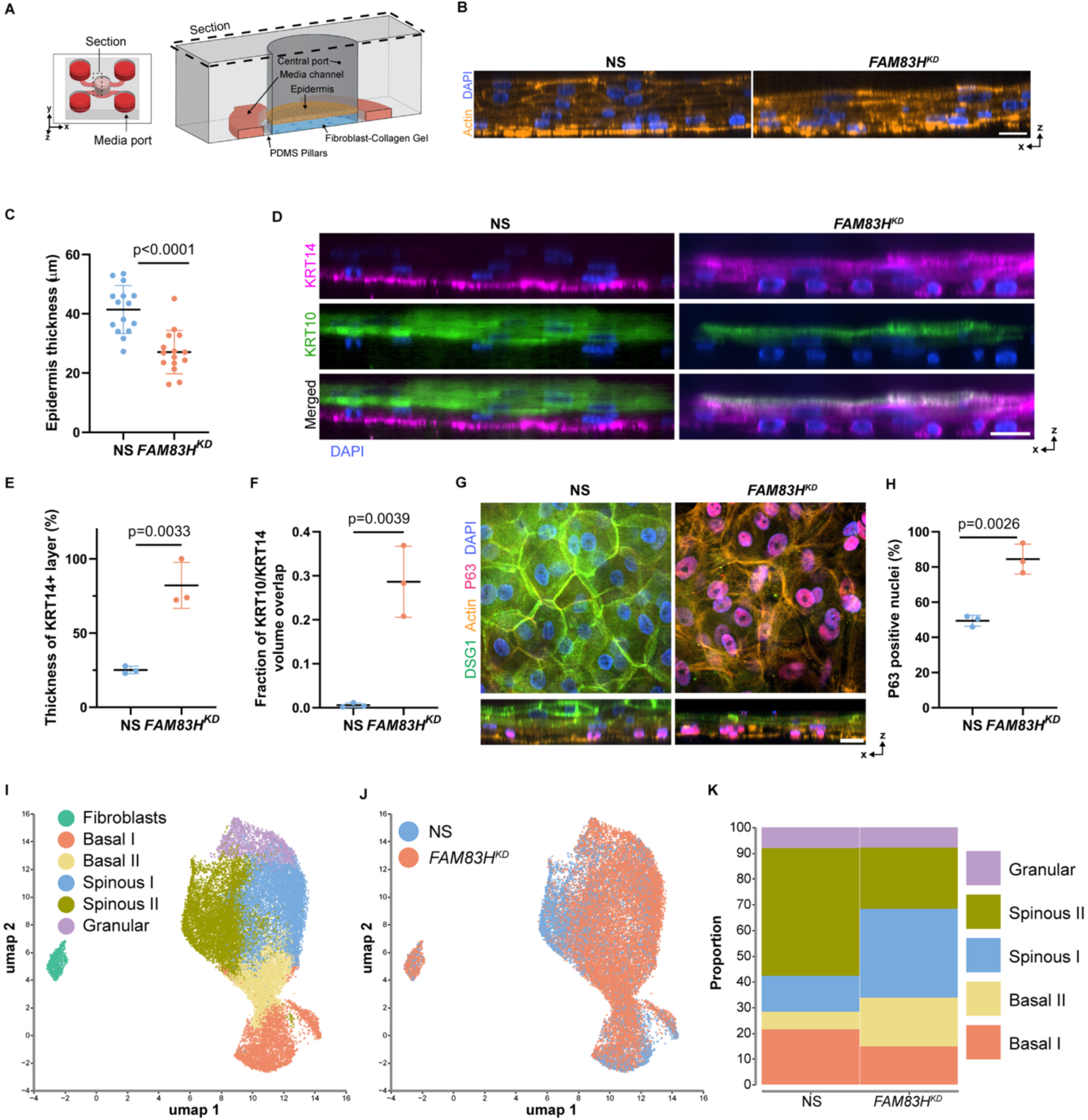
Loss of FAM83H dysregulates basal keratinocyte differentiation and epidermal morphogenesis in a 3D human microphysiological system. A) 3D schematic and cross-sectional view of the StrataChip, highlighting the central port housing the epidermal tissue and dermal equivalent hydrogel, surrounded by media channels. (Adapted from (Amakor et al. 2026)). B) Representative micrographs showing the epidermis in orthogonal slices at day 7, phalloidin stained for F-actin (orange) and DAPI stained for nuclei (blue). Scale bar, 20 μm. C) Quantification of total epidermal thickness in NS and *FAM83H^KD^* 3D epidermal tissues. N = 15 independent devices. D) Orthogonal slice fluorescence micrograph of 3D epidermal tissue at day 7, immunostained with basal and suprabasal layer markers: KRT14 (magenta), KRT10 (green) and DAPI stained for nuclei (blue). Scale bar, 20 μm. E) Quantification of KRT14+layer thickness as a percentage of total epidermal thickness in NS and *FAM83H^KD^* 3D epidermal tissues. F) Quantification of overlap volume of KRT14/KRT10 in NS and *FAM83H^KD^* 3D epidermal tissues. N = 3 independent devices. G) Fluorescence micrographs of the basal layer epidermis from NS and *FAM83H^KD^* 3D epidermal tissues, immunostained for P63, DSG1 and phalloidin stained for F-actin stained and DAPI stained for nuclei. Scale bar, 20 μm. H) Quantification of percentage of P63 positive nuclei NS and *FAM83H^KD^* 3D epidermal tissues. N = 3 independent devices. I) UMAP plot of cell types identified in 3D epidermal tissues. J) UMAP plot of NS and *FAM83H^KD^*cell populations in 3D epidermal tissues. K) Proportion of custom unified two basal, two spinous and one granular layer. In all plots, data represent mean ± SD. Statistical significance was determined by unpaired two-tailed student’s t-test. Exact p values are displayed on each plot; ns, non-significant.

*FAM83H^KD^* resulted in epidermal tissues that multilayered but exhibited reduced tissue thickness compared to NS control (Fig. 2B, C). Interestingly, consistent with our observations in murine models, immunofluorescence staining revealed *FAM83H^KD^* tissues had pronounced expansion of KRT14-positive keratinocytes into the uppermost layers and increased numbers of dual KRT14/KRT10-positive keratinocytes (Fig. 2D-F). To confirm dysregulated expansion of basal keratinocytes, immunostaining for the stemness and self-renewal regulator p63 revealed markedly increased p63-positive nuclei in the *FAM83H^KD^*tissue (Fig. 2G, H). Consistent with murine skin, DSG1 expression was reduced in the suprabasal layer of *FAM83H^KD^* tissues compared to the uniform expression in NS tissues (Fig. 2G). Together, these findings suggest that loss of FAM83H is sufficient to disrupt epidermal fate and mechanics in a 3D human microphysiological model.

To gain deeper insight into how FAM83H alters epidermal development, we performed single-cell RNA sequencing on 3D NS and *FAM83H^KD^* epidermal tissues generated within the StrataChip model. Six epidermal tissues were enzymatically detached from the underlying hydrogel, dissociated, and analyzed. Leiden clustering identified seven distinct cell populations, which were annotated as basal 1 and 2, spinous 1 and 2, granular, and fibroblast cell populations based on established human skin datasets (Fig. 2I, J and Supplementary Fig. 2B, C) (Shuxiong Wang et al. 2020; Zou et al. 2021; Polito et al. 2023; Wiedemann et al. 2023; Amakor et al. 2026). Population proportion analysis corroborated our immunofluorescence findings, revealing an expanded basal cell population in *FAM83H^KD^*3D tissues (Fig. 2K). Specifically, the proportion of basal 2 and spinous 1 subpopulations increased at the expense of basal 1 and spinous 2 subpopulations in *FAM83H^KD^* tissues relative to NS tissues. This finding further supports the hypothesis that the differentiation defect lies at the basal-to-spinous transition (Fig. 2K). Altogether, immunostaining and single-cell transcriptomic analyses of 3D human epidermal tissues demonstrate that FAM83H is essential for early epidermal differentiation and influences basal keratinocyte differentiation and mechanics.

### FAM83H regulates activation of Notch1 signaling during epidermal development

We next analyzed signaling features of the two cell populations (basal 1 and spinous 2) present in NS tissues but reduced in *FAM83H^KD^* samples (Fig. 3A, C). To assess signaling associated with each cluster, we performed Gene Ontology analysis comparing each cluster to the remaining basal and spinous populations, respectively. Several pathways related to basal keratinocyte differentiation were observed (e.g., Hippo and Wnt signaling) with Notch signaling, notably, positively and prominently present in both clusters (Fig. 3B, D).

**Fig. 3.**
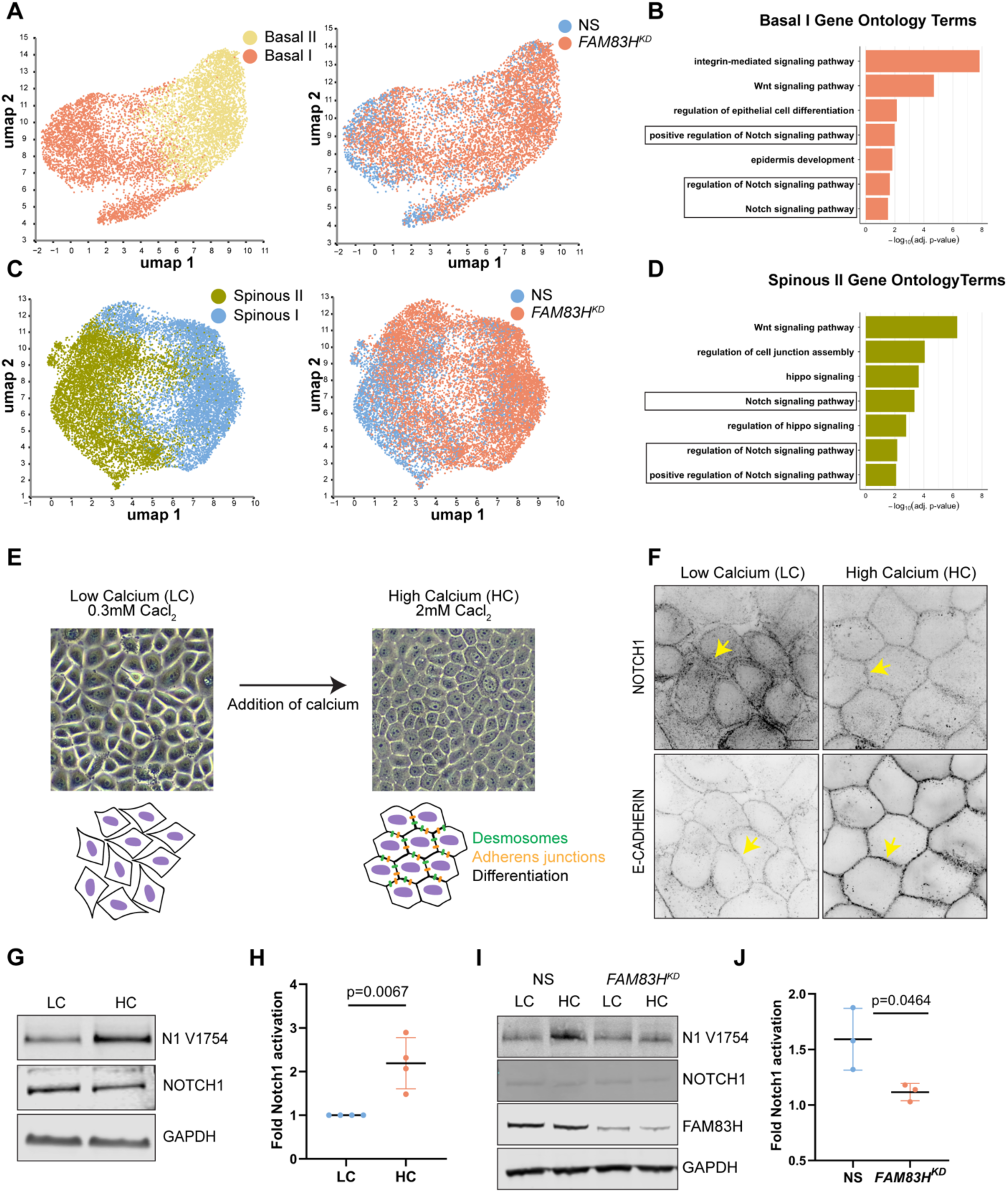
FAM83H regulates activation of Notch1 signaling during epidermal development. A) UMAP of basal cell populations (left) and distribution of NS and *FAM83H^KD^* cells (right). B) Representative Gene Ontology (GO) terms of the distinct basal cell population present in NS condition. C) UMAP of spinous cell populations (left) and distribution of NS and *FAM83H^KD^*cells (right). D) Representative Gene Ontology (GO) terms of the distinct spinous cell population present in NS condition. E) *In vitro* differentiation model of keratinocytes. Bright-field microscopic images of Ker-CT keratinocytes under low and high calcium conditions (left). Illustration of cells under low- and high-calcium conditions (right). F) Representative z-slice micrographs of Ker-CT keratinocytes under low-and-high calcium conditions, immunostained for NOTCH1 and E-cadherin. Scale bar, 10 μm. Yellow arrows highlight the respective staining in some cells. G) Western blot of HaCaT and Ker-CT cell lysates under low and high calcium conditions, immunoblotted for cleaved NOTCH1, total NOTCH1 and GAPDH. H) Quantification of cleaved NOTCH1 signal normalized to GAPDH. N = 4 independent experiments. I) Western blot of the NS and *FAM83H^KD^* Ker-CT cell lysates under low- and high-calcium conditions, immunoblotted for cleaved NOTCH1 (V1754), total NOTCH1, FAM83H and GAPDH. J) Quantification of cleaved NOTCH1 signal normalized to GAPDH. N = 3 independent experiments. In all plots, data represent mean ± SD. Statistical significance was determined by unpaired two-tailed student’s t-test. Exact p values are displayed on each plot; ns, non-significant.

Interestingly, our previous work identified FAM83H as a NOTCH1-interacting protein associated with maintenance of breast epithelial apicobasal polarity and adhesion (White et al. 2023). Notch1 signaling is a critical regulator of epidermal development and homeostasis that promotes basal cell cycle exit and drives differentiation toward the spinous lineage (Rangarajan et al. 2001; Blanpain et al. 2006; Villeneuve et al. 2026). Loss of Notch1 leads to epidermal hyperplasia and tumors resembling basal cell and cutaneous squamous cell carcinomas in mice (Nicolas et al. 2003; Proweller et al. 2006; Demehri et al. 2009). Given Notch signaling was a positive GO feature of cell clusters reduced in *FAM83H^KD^* samples, we next asked whether FAM83H controls Notch1 receptor activation during epidermal keratinocyte differentiation. To test this hypothesis, we employed *in vitro* keratinocyte differentiation assay upon calcium induction (Hennings et al. 1980; Deyrieux and Wilson 2007) (Fig. 3E). Calcium promotes the formation of desmosome cell-cell junctions and initiates the differentiation of keratinocytes toward suprabasal lineages (Micallef et al. 2009; Bikle et al. 2012). Indeed, immunofluorescence staining of keratinocytes cultured under low-versus-high calcium showed a marked junctional localization of E-cadherin along with concomitant enrichment of NOTCH1 at cell-cell junctions upon calcium induction (Fig. 3F), and this NOTCH1 junctional localization is unchanged in the absence of FAM83H (Supplementary Fig. 3F). NOTCH1 is a transmembrane receptor that is proteolytically activated to exert transcriptional signaling (Kovall et al. 2017). To assess Notch1 receptor proteolytic activation, we performed immunoblot analysis using a cleavage-specific NOTCH1 V1754 antibody that recognizes the activated form of the receptor (Polacheck et al. 2017; White et al. 2023). We observed robust proteolytic Notch1 activation upon calcium induction (Fig. 3G, H). Consistent with a role in regulating Notch1 signaling during keratinocyte differentiation, *FAM83H^KD^* cells significantly reduce Notch1 proteolytic activation upon calcium induction (Fig. 3I, J). Together, single-cell transcriptomic analyses of 3D epidermal tissues and *in vitro* biochemical assays indicate that FAM83H is required for proper Notch1 proteolytic activation and signaling during keratinocyte differentiation.

### FAM83H complexes Notch1 with DSP and keratin to control receptor activation

Keratin network organization is the sole prescribed cellular function of FAM83H (Kuga et al. 2013, 2016; Tokuchi et al. 2021). We therefore investigated whether this function was related to its regulation of Notch1 activation. We first characterized FAM83H and keratin expression in keratinocytes and examined the effects of FAM83H loss on keratin filament organization and desmosomes. Immunostaining confirmed that FAM83H co-localizes with the keratin filament network at cell-cell junctions and throughout the cytoplasm (Supplementary Fig. 3A). We generated CRISPR-mediated FAM83H knockout keratinocytes (*FAM83H^KO^)*. Upon loss of FAM83H, the keratin filament network collapsed to a perinuclear locus, with markedly fewer filaments extending to the cell periphery compared to the extensive peripheral network in control cells (Fig 4A). Quantification using the coefficient of variation of keratin signal intensity demonstrated increased spatial heterogeneity and disrupted filament organization in *FAM83H^KO^* cells (Fig. 4B). To determine whether filament collapse was spatially linked to the microtubule organizing center (MTOC), we co-stained for α- and γ-tubulin; collapsed keratin aggregates co-localized with α-tubulin-enriched regions and were associated with the MTOC marked by γ-tubulin, suggesting that the loss of FAM83H causes keratin filaments to retract retrogradely and accumulate at the MTOC (Supplementary Fig. 3B, C). We next examined desmosomes by immunostaining for DSP, a core desmosomal plaque component. In control keratinocytes, DSP localized in a tight, continuous linear pattern along cell-cell borders, whereas *FAM83H^KO^* cells displayed a discontinuous, punctate, and misoriented distribution, indicating impaired desmosome assembly or stability at the membrane (Fig. 4A). Consistent with desmosome dysregulation, *FAM83H^KO^* cells exhibited reduced cell-cell cohesion in a dispase-based dissociation assay, evidenced by increased sheet fragmentation under mechanical stress (Supplementary Fig. 3D, E). Together, these data extend prior reports and define a FAM83H loss phenotype characterized by perinuclear keratin collapse at the MTOC, desmosome disorganization, and reduced intercellular cohesion, raising the possibility that cytoskeletal disruption may impair the cortical membrane dynamics required for Notch1 activation.

**Fig 4.**
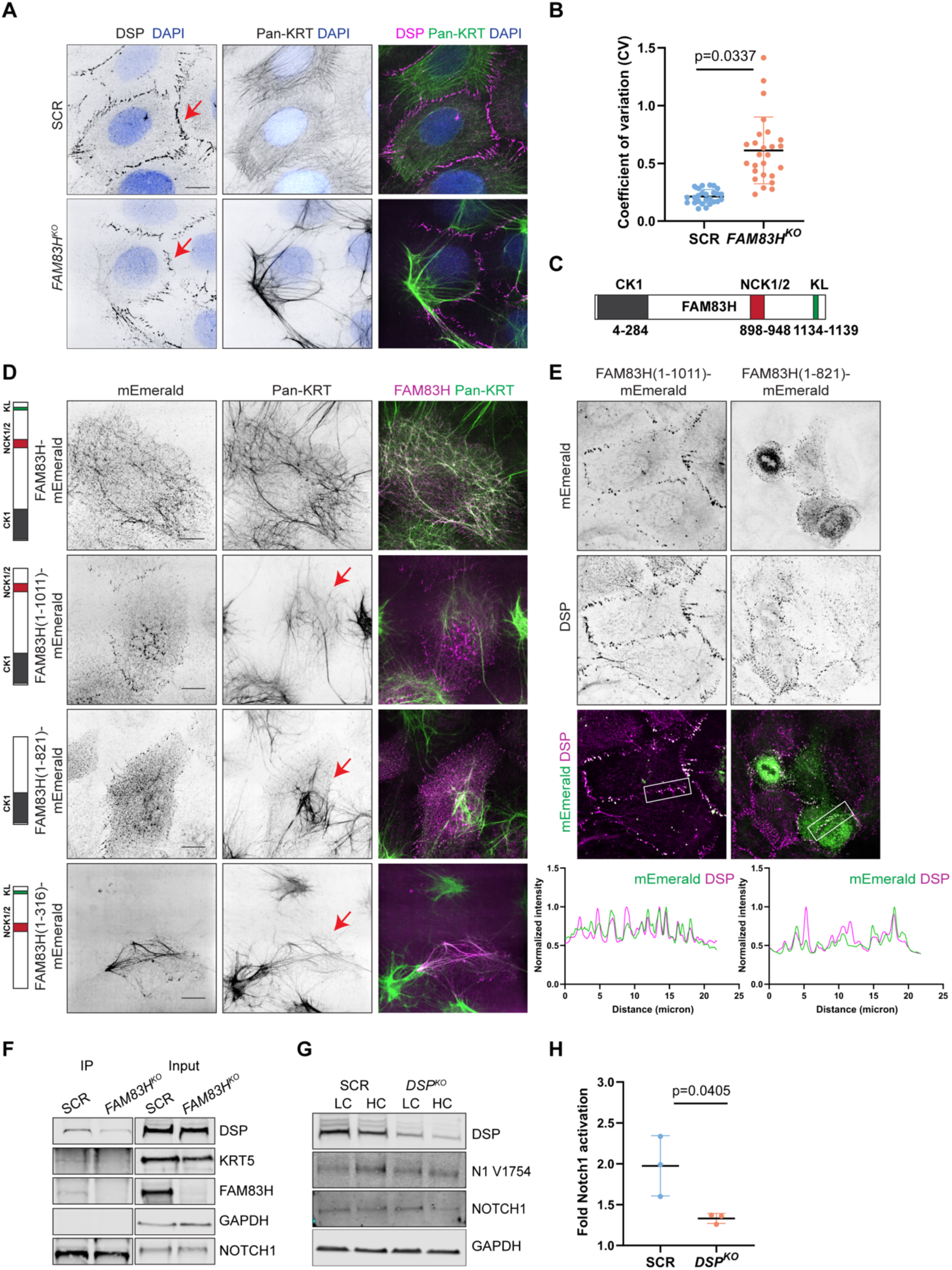
FAM83H complexes Notch1 with DSP and keratin to control receptor activation. A) Representative z-slice micrographs of SCR and *FAM83H^KO^* HaCaT cells immunostained for Pan-KRT and DSP and merge of Pan-KRT (green) and DSP (magenta). Red arrows indicate DSP pattern. Scale bar, 10 μm. B) Coefficient of variation (CV = SD/mean) was calculated for individual cells as a measure of keratin distribution uniformity. Each dot represents a single-cell measurement pooled from 3 independent experiments. Statistical analysis was performed on replicate means using an unpaired two-tailed student’s t-test. C) Schematic of FAM83H full-length protein with three known interaction domains. CK1 binding domain in N-terminus, NCK1/2 binding domain and KL, keratin localization in C-terminus. D) Representative z-slice micrographs of *FAM83H^KO^* HaCaT keratinocytes transfected with full length and truncated constructs of FAM83H-Emerald and immunostained for Pan-KRT. Merged images with KRT (green) and FAM83H (magenta). Red arrow indicates the keratin disorganization. Scale bar, 10 μm. E) Representative z-slice micrographs of *FAM83H^KO^* HaCaT keratinocytes transfected with FAM83H-Emerald truncated constructs 1-1011 and 1-821 and immunostained for DSP and merged images with DSP (magenta) and Emerald (green). Scale bar, 10 μm. Line-scan analysis of Emerald and DSP fluorescence intensity indicated in the above images (normalized to respective maximum intensity). F) Western blot of the SCR and *FAM83H^KO^* HaCaT cell lysates followed by the NOTCH1 immunoprecipitation, immunoblotted for DSP, KRT5, FAM83H, GAPDH and NOTCH1. G) Western blot of the cell lysates from SCR and *DSP^KO^* Ker-CT keratinocytes under low- and high-calcium conditions, immunoblotted for cleaved NOTCH1 (V1754), total NOTCH1, DSP and GAPDH. H) Quantification of NOTCH1 activation normalized to GAPDH. N = 3 independent experiments. In all plots, data represent mean ± SD. Statistical significance was determined by unpaired two-tailed student’s t-test. Exact p values are displayed on each plot; ns, non-significant.

To investigate domain-specific functions of FAM83H, we designed a series of truncation constructs guided by AlphaFold structural predictions and ConSurf sequence conservation analysis, ensuring that no truncations fell within predicted structural domains or highly conserved regions. We then assessed the localization of each construct and its ability to rescue keratin disorganization in FAM83H-depleted keratinocytes. The FAM83H protein is largely intrinsically disordered but contains three characterized binding sites located in its N- and C-terminal regions, which mediate interactions with CK1, NCK1/2, and keratin filaments (Fig. 4C) (Tachie-Menson et al. 2020; Kuga et al. 2022). We generated three FAM83H truncated constructs lacking the CK1 binding, NCK1/2 binding, and keratin localization domains (residues 316-1179, 1-821 and 1-1011 respectively). When expressed in CRISPR-mediated *FAM83H^KO^* HaCaT keratinocytes, only the full-length construct rescued keratin disorganization (Fig. 4D). Although none of the truncations restored keratin organization, the 1-821 and 1-1011 constructs retained junctional localization and co-localized with DSP (Fig. 4E), indicating that FAM83H has junction-targeting motifs independent of its ability to bind keratin. The 316-1179 truncation, which lacks N-terminal CK1-binding domain, co-localized with keratin filaments, but was unable to rescue the keratin disorganization.

Given FAM83H localizes to desmosomes and modulates keratin and desmosome organization, we next examined whether these functions are associated with the NOTCH1 regulation. We performed co-immunoprecipitation and observed that NOTCH1 interacts with FAM83H, KRT5, and DSP in keratinocytes under both low- and high-calcium conditions (Supplementary Fig 3G). Interestingly, co-immunoprecipitation of NOTCH1 in SCR and *FAM83H^KO^* keratinocytes revealed diminished binding of NOTCH1 with KRT5 and DSP in the absence of FAM83H (Fig. 4F), indicating that NOTCH1 association with KRT5 and DSP is dependent on FAM83H. To assess whether this interaction was critical to Notch1 activation, we generated CRISPR-mediated DSP knockout keratinocytes (*DSP^KO^*). *DSP^KO^*cells had decreased Notch1 proteolytic activation in the *in vitro* calcium induction assay (Fig 4G, H). Collectively, these data support a model in which FAM83H, keratin, and NOTCH1 form a protein complex at the junction, and this complex is required for Notch1 activation to promote epidermal differentiation.

## Discussion

Our study identifies FAM83H as a previously unrecognized regulator of epidermal morphogenesis that mechanistically links keratin cytoskeletal organization to Notch1 signaling and keratinocyte differentiation. Using complementary *in vivo* mouse model and a human 3D microphysiological epidermal system, we demonstrate that FAM83H loss disrupts epidermal architecture by impairing basal cell differentiation and cell-cell cohesion. Mechanistically, our data support a model in which FAM83H organizes keratin filaments at cell-cell junctions to promote the assembly of a DSP-keratin-Notch1 complex required for efficient Notch1 proteolytic activation and the basal-to-spinous transition during epidermal morphogenesis.

FAM83H is broadly expressed across epidermal layers and is essential for proper epidermal development and homeostasis. The cell shape, adhesion, and keratin organization defects observed in *Fam83h^-/-^* mice are consistent with a cell-autonomous role for FAM83H in maintaining epidermal architecture, though the structural basis of these defects *in vivo* remains to be fully defined. Future studies employing higher-resolution approaches, such as electron microscopy, will be important to characterize ultrastructural changes at desmosomes and the keratin cytoskeleton in *Fam83h^-/-^* epidermis. Although FAM83H mutations cause amelogenesis imperfecta in humans, overt skin disease has not been reported in affected patients. Interestingly, *FAM83H* knockout mice exhibit skin abnormalities but do not develop enamel defects, whereas disease-causing truncation mutations reproduce enamel defects in mice, suggesting neomorphic effects of mutant FAM83H during enamel development (Shih-Kai Wang et al. 2015; S.-K. Wang et al. 2019). In addition, embryonic lethality observed in *Fam83h*^⁻/⁻^ mice suggests that complete loss of FAM83H function may be incompatible with development, which may explain why only truncation mutations have been identified in humans (S. K. Wang et al. 2021).

Recapitulation of key *Fam83h*^⁻/⁻^ phenotypes, including basal cell expansion, impaired differentiation, and desmosome defects, in our human 3D microphysiological model supports a conserved and keratinocyte-intrinsic role for FAM83H across species. Because this system lacks immune cells and melanocytes, the phenotypes observed can be attributed to cell-autonomous keratinocyte mechanisms, strengthening mechanistic interpretation. Incorporating additional cell types in future adaptations of this model will be important for capturing the full complexity of FAM83H function at the tissue level.

Single-cell transcriptomic analysis revealed that the cell populations most reduced by FAM83H depletion, Basal I and Spinous II, are enriched for Notch1 signaling. This is consistent with the established role of Notch1 as a direct driver of basal cell cycle exit and commitment to the spinous lineage (Rangarajan et al. 2001; Blanpain et al. 2006), and with the observation that Notch1 loss in mice produces epidermal hyperplasia and tumors resembling basal and cutaneous squamous cell carcinomas (Nicolas et al. 2003; Proweller et al. 2006; Demehri et al. 2009). The expansion of dual KRT14/KRT10-positive cells in *Fam83h*^⁻/⁻^ epidermis closely phenocopies impaired Notch1 signaling, supporting the notion that FAM83H-dependent Notch1 activation is critical during early epidermal development. Interestingly, Notch1 junctional localization and expression are maintained in *Fam83h*^⁻/⁻^ mice at postnatal stages examined (Supplementary Fig. 3H), suggesting that the requirement for FAM83H in Notch1 activation may be most acute during earlier developmental windows. Consistent with this interpretation, a recent study demonstrated that Notch1 is activated in a subset of crowded basal cells to facilitate their delamination and differentiation (Villeneuve et al. 2026), a process that may be particularly sensitive to FAM83H-dependent keratin organization.

Our domain truncation analysis reveals that FAM83H harbors functionally distinct domains that independently regulate keratin cytoskeletal organization and junctional localization. Although truncations lacking the keratin localization domain failed to rescue keratin disorganization, they retained the ability to localize to cell-cell junctions and co-localize with DSP. This indicates that FAM83H contains junction-targeting sequences that are separable from its keratin-binding activity and suggests that FAM83H coordinates multiple structural and signaling functions at epithelial junctions through modular domain organization. Our co-immunoprecipitation data demonstrate that Notch1 associates with KRT5 and DSP in a FAM83H-dependent manner, and that DSP loss alone is sufficient to impair Notch1 proteolytic activation. These findings extend an emerging paradigm in which keratin intermediate filaments directly participate in cell signaling. In colonic epithelium, KRT8 directly interacts with Notch1 to regulate differentiation and cell fate, and KRT8 deletion reduces Notch1 levels and activity (Lähdeniemi et al. 2017); our data suggest an analogous mechanism operates in epidermal keratinocytes, where KRT5 participates in a FAM83H-organized junctional complex that licenses Notch1 activation. Notably, single-cell sequencing also identified enrichment of Hippo and Wnt signaling in the cell populations lost upon FAM83H depletion. Given the broad keratin disorganization caused by FAM83H loss, and the established roles of keratins in modulating these pathways, including KRT14-dependent regulation of Hippo signaling during early differentiation (Guo et al. 2020), it is plausible that FAM83H loss broadly perturbs multiple signaling axes. Systematically defining these additional FAM83H-dependent signaling relationships will be an important direction for future work.

An outstanding mechanistic question is how the FAM83H-DSP-keratin-Notch1 complex promotes receptor proteolytic cleavage; future studies should address whether this complex influences γ-secretase accessibility or mechanical forces required for Notch1 proteolytic activation. Our findings have potential implications for epithelial disease. FAM83H expression is reduced in cutaneous squamous cell carcinoma and basal cell carcinoma relative to normal epidermis (Tokuchi et al. 2021) and given the established tumor-suppressive function of Notch1 signaling in skin, the ability of FAM83H to sustain Notch1 activation provides a plausible mechanism by which FAM83H downregulation could contribute to epidermal tumorigenesis. Also, loss-of-function DSP mutations cause skin fragility disorders, raising the possibility that impaired Notch1 signaling may contribute to these conditions (Armstrong et al. 1999; Norgett et al. 2000; Nitoiu et al. 2014). More broadly, our work establishes FAM83H as a molecular scaffold that coordinates keratin cytoskeletal architecture with junctional signaling during epidermal morphogenesis, and positions keratin-associated proteins as active participants in the epithelial fate decisions that govern tissue homeostasis and disease.

## Declaration of Interests

The authors declare no competing interests

## Materials and methods

### Animals

Animals were housed in individually ventilated cages (Tecniplast) with free access to water and food (Altromin), Safe Select Fine bedding (Velaz), and environmental enrichment under a 12/12 light-dark cycle. The study complied with EU laws (Project licence AVCR 4212/2023 SOV II). Experiments followed the Czech Centre for Phenogenomics (CCP) Standard Operating Procedures. Fam83h knockout mice (Fam83hem2(IMPC)Ccpcz, shortly, *Fam83h^-/-^)* were generated using CRISPR/Cas9 technology within the IMPC effort (www.mousephenotype.org) as described in (Ogan et al. 2026).

### Cryopreservation and Sectioning of mouse skin tissue

Dorsal skin samples were collected from mice at early (2-3 weeks) and adult (16 weeks) time points. Prior to tissue collection, hair was carefully trimmed from the dorsal region. Skin specimens were fixed overnight at room temperature in 4% formaldehyde (FA), followed by transfer to 70% ethanol for storage at 4°C until processing. Back skin tissues from *Fam83h^+/+^* and *Fam83h^-/-^* mice were first fixed in FA and shipped to UCSF in phosphate-buffered saline (PBS). Upon arrival, tissues were left in 30% sucrose in PBS overnight at 4°C. They were then embedded in optimal cutting temperature (OCT) compound, frozen and stored at -80°C until use. OCT-embedded skin tissues were sectioned at a thickness of 20 µm using a cryostat (Microm HM525NX). Sections were air-dried overnight and either processed immediately for immunostaining or stored at -20°C for later use.

### EdU incorporation assay

Mice were injected intraperitoneally with EdU working solution at a dose of 150 μg EdU in 100 μL PBS per animal. The working solution was prepared by diluting the 10 mM EdU stock 1.6-fold in PBS to a final concentration of 6.25 mM. Animals were sacrificed 3 h after injection, and skin tissues were dissected and fixed in 4% formaldehyde in PBS overnight.

### H&E Staining

Tissues samples were fixed in 10% buffered neutral formalin 24 hours, processed by Leica ASP6025 automated vacuum processor, embedded in Paraplast X-tra (Leica Biosystems) manualy using paraffin-embedding station Leica EG1150 H+C. Formalin-Fixed Paraffin-Embedded (FFPE) blocks were serially sectioned on Leica Fully Motorized Rotary Microtome RM2255-FU at 4,5 μm thickness. Slides were deparaffinized, rehydrated through graded ethanol, and stained by Harris hematoxylin (Biognost) for nuclear visualization and Eosin Y (Carl Roth) for cytoplasmic and extracellular matrix visualization, dehydrated using an automated staining and coverslipping system (Leica ST5010-CV5030 Stainer Integrated Workstation with automatic coverslipping, Leica Biosystems). Images were acquired Leica DM3000 LED microscope with a 20x Objective (10x zoom) and analyzed in Leica Application Suite V4.13 software.

### Immunofluorescence of mice skin sections

Slides were removed from -80°C, allowed to sit at RT for 10-15 min. Sections were bordered using PapPen to prevent cross-contamination of antibodies. Sections were permeabilized with PBS-Triton X-100 (0.1%) and washed 2X with PBS for 5 min each wash. Blocking solution was prepared fresh with the following final composition 1% BSA, 0.01% Gelatin, 2.5% Goat serum, 0.3% Triton X with 1X PBS using stock solution of 10% BSA, 1% Gelatin, Goat serum and 10% Triton X. Sections were blocked for 1 hr. Sections with primary antibodies were incubated for 2 hr at RT and washed 3X with PBST for 5 min each wash. Further incubated with secondary antibodies (1:400), rhodamine phalloidin (1:2000) and DAPT (1:1000) for 1 hr at RT and washed 3X with PBST for 5 min each wash. Tissue sections were sealed with cover slips using Glass Antifade Mountant for imaging.

### EdU staining

Edu detection was performed according to the Invitrogen Click-iT EdU Kit protocol. Skin sections were incubated with the EdU mix for 30 min and followed by DAPI and phalloidin dyes for 30 min. Sections were then sealed with coverslips on top using Glass Antifade Mountant (Invitrogen ProLong) for imaging.

### Cell lines and culture conditions

Ker-CTs (ATCC) were maintained with KBM-Gold medium (Lonza) supplemented with KGM Gold keratinocyte growth medium BulletKit (Lonza). For high-calcium condition, 1.7 mM calcium chloride was added to the media and for low calcium condition, regular media was used. Neonatal Human Dermal Fibroblasts (NHDFs) (Lonza) were maintained with high glucose DMEM (Gibco) supplemented with 10% fetal bovine serum (Peak or SeraPrime), and 1% penicillin/streptomycin (Sigma). HaCaT cells (provided by Wittmann lab, UCSF) were maintained in DMEM (Gibco) medium consisting of 100U/ml penicillin, 100 μg/ml streptomycin (Invitrogen) and 10% Fetal bovine serum (FBS). To prepare no calcium media DMEM no calcium (Invitrogen) was used and FBS was calcium depleted by incubating with chelating resin mesh (Bio-rad) for 1 hr at 4°C. Calcium chloride was added to the media to make low- and high-calcium conditions (0.03 mM and 1.5-2 mM, respectively). For rescue experiments, *FAM83H^KO^* cells were transfected with full-length and truncated FAM83H constructs using JETOPTIMUS or transduced via lentiviral delivery. All cells were routinely tested for *Mycoplasma* via PCR test.

### Antibodies and reagents

Antibodies against Notch1 ICD (D1E11, 1:100 IF, 1:1,000 WB); Notch1 V1744 (D3B8, 1:500 WB); GAPDH (14C10, 1:10,000 WB); KRT5 (D4U8Q, 1:1000 WB, 1:100 IF); and P63 (D9L7L, 1:1000 WB) were from Cell Signaling Technologies; Cyto Pan-KRT (1:200 IF); DSG1 (1:100 IF); and γ Tubulin Antibody (D-10 1:100 IF) from SCBT; E-cadherin antibody (HECD-1, 1:1,000 IF, 1:1,000 WB) was from Takara Bio; FAM83H antibodies 328 and 327 (1:100 IF 1:1,000 WB); Desmoplakin (1:100 IF 1:2000 WB) were from Bethyl Laboratories; KRT14 (PIPA516722, 1:100 IF); KRT10 (MA106319, conjugated with Alexa fluor 488 1:25 IF); Rhodamine phalloidin (1:2000); and Alexa Fluor 488, 568, and 647 goat anti-mouse and anti-rabbit IgG secondary antibodies (1:400) were from Invitrogen.

### Lentiviral-mediated gene editing

Stable CRISPR-modified HaCaT and Ker-CT cell lines were generated using the lentiCRISPRv2 system following our previously established protocols (Polacheck et al. 2017). Specific guide RNAs were cloned into the BsmBI site of lentiCRISPRv2: SCR, 5′-GTATTACTGATATTGGTGGG-3′; *NOTCH1*^KO^, 5′-CGTCAGCGTGAGCAGGTCGC-3′; *FAM83HKO*, 5′-CTGTTCGAGAAGCTTCGCGG-3′; and *DSPKO*, 5′-GAGATGGAATACAACTGACT-3′.

ShRNA-mediated HaCaT and Ker-CT cell lines were generated by cloning shRNA sequences into modified pLL3.7 vectors using T4 DNA Ligase (New England Biolabs; M0202L). A non-targeting control NS (5′-ATCGACTTACGACGTTAT-3′) and an shRNA targeting *FAM83H* (5′-GGAACTGATTTAAGAAACA -3′) were used.

sgRNA-containing lentiCRISPRv2 plasmids and shRNA-containing pLL3.7 plasmids were co-transfected with psPAX2 (plasmid #12260; Addgene) and pMD2.G (plasmid #12259; Addgene) packaging plasmids into HEK293T cells using calcium phosphate transfection. After 48 hr, viral supernatants were collected, concentrated using 4× lentivirus concentrator (PEG-IT), and resuspended in PBS. HaCaT and Ker-CT cells were transduced in their corresponding growth medium overnight and given fresh medium the following morning. At 48 hr after transduction, cells were passaged and plated in 6-well plates at 1.5 × 10^5^ cells per well and selected with 2 μg/ mL puromycin for 4 days. Gene knockdown and knockout efficiency were verified by Western blot.

### Transient Transfection

HaCaT cells were seeded in 12-well plates and transfected with 0.5-2 μg plasmid DNA using jetOPTIMUS® DNA transfection reagent (Polyplus, VWR) according to the manufacturer’s instructions. Plasmid DNA was mixed with jetOPTIMUS reagent in jetOPTIMUS buffer and incubated for 10 min at room temperature. Media was replaced 4-6 h post-transfection, and cells were analyzed 24-96 h later.

### Microfluidic device design and fabrication

#### Device design

The device was designed as described in (Amakor et al. 2026). Device consists of 4 total media ports and a central port surrounded by PDMS pillars that are each 400 μm tall to contain the dermal equivalent. Each media port pair is connected with a channel that runs from one media port, alongside the central port and then to the other media port.

#### Device fabrication

The device was fabricated as described in (Amakor et al. 2026; Polacheck et al. 2019). In brief, silicon master was fabricated using photolithography as described in (Polacheck et al. 2019) using the Alveole PRIMO system. Devices were then made through soft lithography with polydimethylsiloxane (PDMS Sylgard 184, Dow Corning) mixed with Sylgard 184 silicone elastomer curing agent (Dow Corning) at a 10:1 ratio by weight. PDMS devices were cut, holed punched, and then surface activated in a plasma cleaner (Harris Plasma) for 30 s at 0.3 Torr. Devices were then bonded to a coverslip and placed in a 100°C oven for 30 min. Devices central port were then treated with a 2 mg/ml solution of dopamine hydrochloride (Sigma) or polydopamine (PDA) was made with 10 mM Tris at a pH of 8.5 for 90 min. Devices were then washed and sterilized in 70% ethanol for 1 hr on a laboratory shaker.

### Engineered 3D epidermal tissue culture

Epidermal tissue model was cultured as described in (Amakor et al. 2026). Devices were UV sterilized for 15 min before the addition of a 3 mg/ml concentration of rat tail collagen 1 (corning) with normal human fibroblasts (NHDFs) a final concentration of 6×10^4^ cells/mL into the central port. Collagen-firboblast gel was then polymerized in an incubator at 37°C for 20 min. NHDF media was then added to the devices and incubated at 37°C. Keratinocytes were then seeded the following day at a concentration of 2.25×10^6^ cells/mL into the central port. Devices were then cultured and stained as described in (Amakor et al. 2026).

### Epidermal device immunofluorescence

All incubation and washing steps in the device were done on a laboratory rocker (VWR) at room temperature unless otherwise specified. For all steps, unless otherwise stated, when all ports were filled, reagents were added to the central port first before the media ports to prevent hydrostatic pressure on the model epidermis. Media was removed from the media ports and devices were fixed by filling all ports with 4% paraformaldehyde (Electron Microscopy Sciences) prepared in 1X PBS with calcium and magnesium (0.9 mM CaCl_2_ and 0.49 mM MgCl_2_) pre-warmed to 37°C. Devices were fixed for 35 min. Devices (all ports) were then washed 3 times for 15 min each with 1X PBS and then quenched with 100 mM glycine (Sigma) in H_2_O for 1 hr. Devices were then washed again as described above and then permeabilized with 0.5% Triton-X (Sigma) in 1X PBS for 90 min. Devices were then washed as described above and then blocked with 10% normal goat serum (Millipore) in 1X PBS blocking buffer for 1 hr. Primary antibodies were prepared in blocking buffer at concentrations detailed in the “Antibodies and reagents” section. Blocking buffer was removed from all ports of the device and then 5 μl of blocking buffer was added only to the central port of each device. 100 μl of diluted primary antibodies was added into each media port pair (200 μl total per device) and incubated overnight on a rocker at 4°C. Devices (all ports) were washed 4 times for at least 15 min each with 1X PBS. Secondary antibodies and fluorescent dyes were prepared in blocking buffer at concentrations detailed in the “Antibodies and reagents” section. Devices were incubated in secondary antibodies and fluorescent dyes as earlier described with the primary antibodies starting with the addition of 5 μl of blocking buffer to the central port and were incubated overnight on a rocker at 4°C protected from light. Devices (all ports) were washed 4 times for at least 15 min each with 1X PBS. Each device was filled with 1X PBS + 1% penicillin/streptomycin (Sigma) and then stored in parafilmed dishes at 4°C until imaged.

### Epidermal model tissue dissociation

Seven devices for each condition were enzymatically dissociated as described in (Amakor et al. 2026). In brief, epidermis was first digested within a dispase digestion mix pre-warmed to 37°C for 90 min with mechanical agitation on a ThermoMixer set to 1000 rpm. After digestion, the solution was spun down at 300xG at RT for 10 min. Pellet was resuspended in a collagenase digestion mix and incubated for 40 min at 37°C with mechanical agitation on a ThermoMixer set to 1000 rpm. Solution was then spun down at 300xG at RT for 10 min then resuspended and incubated in 500 µl of 0.25% trypsin-EDTA for 10 min at 37°C in the ThermoMixer at 1000 rpm. Digestion solution was then neutralized with 1000 µl of DMEM media and filtered through a 70 µm cell strainer. Filtered solution was finally spun down at 300xG at RT for 10 min and then resuspended in 50 µl 0.2% BSA and kept on ice until cell fixation.

### Single-Cell RNA sequencing analysis

Dissociated model epidermis cells were fixed, sequenced and processed as described in (Amakor et al. 2026). Data processing was done largely as in (Amakor et al. 2026) with minor modifications specifically relevant for this dataset. The dataset used here underwent Trailmaker’s automatic quality control and filtering to remove cells with low transcripts (<1247.548), high percentage of mitochondrial reads (>24.56%), and high doublet probability score (>0.79) from further downstream analysis. After removal of these cells, 19,293 cells remained with a median number of 7334 transcripts per cell. Data were normalized using log-normalization and subsequently integrated using the Fast MNN method with the number of highly variable genes (HVG) and principal components set to 2000 and 30, respectively. Graph-based clustering was performed with the Leiden algorithm at a resolution of 0.36. Clusters were then visualized on Uniform Manifold Approximation and Projection (UMAP) embedding. Automated Leiden clusters were manually annotated based on expression profiles of known epidermal cell fate markers such as, but not limited to, KRT14, KRT10, Involucrin, Loricrin, and previously identified epidermal cell populations (Shuxiong Wang et al. 2020; Zou et al. 2021; Polito et al. 2023; Wiedemann et al. 2023). UMAP and dot plots were generated on Trailmaker. DEGs and Gene ontology analysis was performed as described in (Amakor et al. 2026): Differentially expressed genes (DEGs) were identified for each cluster of interest compared to the rest of the cell sample with the presto implementation of the Wilcoxon rank sum test and auROC analysis. Generated DEGs were then filtered on R studio for genes with log fold change (logFC) > 0.25 and adjusted p-value < 0.05. Gene Ontology (GO) analysis of DEGs was done on R studio using the clusterProfiler R package and the Bioconductor homosapien genome wide annotation database (‘org.Hs.eg.db’). GO analysis was then visualized using the ggplot2 R package. Representative terms for each cluster were then selected from generated GO terms. GO terms are listed in Supplementary Table 1 and 2.

### Immunofluorescence of 2D monoloayers

For immunofluorescence of 2D monolayers, cells (HaCaT and Ker-CT) were plated on 18-mm glass coverslips coated with collagen. Coverslips were surface activated by plasma treatment for 30 s, APTES for 30 min and glutaraldehyde for 1 hr, and washed three times with dd water. Coverslips were stored at RT for maximum 1 month. 50μg/ml collagen type I in PBS was added to the coverslip and spread around the surface and incubated at 37°C for 45 min. Cells were resuspended in their corresponding culture medium and 1 × 10^5^ cells were added to each well. Once cells reached a confluent monolayer, coverslips were fixed in 4% paraformaldehyde in PBS++ for 15 min at 37°C, rinsed three times with PBS, and permeabilized in 0.1% Triton X-100 for 10 min. For methanol fixation and permeabilization, coverslips were submerged in ice-cold methanol for 5 min at 20°C. Coverslips were washed three times with PBS and blocked in 10% goat serum in PBS for 1 hr. Primary and secondary antibodies were applied in 10% goat serum in PBS for 1 hr at room temperature and rinsed three times with a ten min interval. Coverslips were sealed on a glass slides using Glass Antifade Mountant for imaging.

### Cloning of FAM83H-mEmerald constructs

Full-length human FAM83H cDNA was amplified by PCR from an MCF10A cDNA library generated by reverse transcription, using primers containing overhangs designed for Gibson assembly into a modified pRRL lentiviral expression vector to generate a FAM83H-mEmerald fusion construct, which was verified by whole-plasmid sequencing. Truncated constructs (residues 316-1179, 1-1011, and 1-821) were generated by PCR amplification from the full-length FAM83H-mEmerald construct and assembled using the Gibson Assembly method. All primers were synthesized by IDT and primer sequences are provided in Supplementary Table 3. Site-directed mutagenesis was performed to generate CRISPR-resistant rescue constructs.

### Immunoblotting

Cells (Ker-CT and HaCaT) were seeded into individual wells of a 6-well plate at a density of 4 × 10⁵ cells per well in low-calcium assay medium and cultured for 48 hr. For high-calcium conditions, media was supplemented with 1.7 mM CaCl₂, while low-calcium media was maintained for control conditions. Cells were grown into confluence and incubated for 6-8 hr at respective calcium conditions before lysing. Monolayers were rinsed with PBS and lysed in buffer containing 50 mM Tris-HCl (pH 7.4), 150 mM NaCl, 1% Triton X-100, 0.1% SDS, 0.5% sodium deoxycholate, and 1.5× protease inhibitor cocktail (Thermo Fisher Scientific). Lysates were sheared by passing through a 21G syringe 10 times, incubated on ice for 10-15 min, and centrifuged at 13,000 × g for 10 min at 4°C. Protein concentrations were determined using a BCA assay (Prometheus), and samples were denatured in 1× NuPAGE LDS sample buffer (Life Technologies) containing 5% β-mercaptoethanol. Equal amounts of protein were resolved by SDS-PAGE and transferred onto nitrocellulose membranes using a Mini Trans-Blot Cell (Bio-Rad). Membranes were blocked in 5% non-fat milk in TBST (0.1% Tween-20) for 1 hr at room temperature and incubated with primary antibodies overnight at 4°C. After washing three times over 30 min with TBST, membranes were incubated with IR Dye-conjugated donkey anti-rabbit or anti-mouse secondary antibodies (1:10,000; LI-COR) for 1 hr at room temperature. Membranes were washed again with TBST over 30 min and imaged using an Odyssey CLx imaging system (LI-COR). Band intensities were quantified using ImageJ. Brightness and contrast adjustments were applied uniformly, and protein levels were normalized to GAPDH unless otherwise specified. Uncropped blots are provided in the source data.

### Dispase Dissociation assay

Confluent HaCaT monolayers in a 6-well plate were washed with prewarmed PBS and incubated with Dispase® II (Sigma) (2.5 U/mL in HBSS; 250 μL per well) for 20 min at 37°C until the cell sheet detached. The dispase solution was removed, and 350 μL of fresh HBSS (Gibco) was added. MTT solution (VWR) (20 μL, 5 mg/mL) was added, and cells were incubated for an additional 10 min at 37°C. Detached monolayers were transferred to microcentrifuge tubes and subjected to mechanical stress by vortexing for 1 min. Cell-cell cohesion was assessed by quantifying the number of fragments generated.

### Microscopy

For microfluidic device imaging, confocal microscopy was conducted on a Yokogawa CSU-W1/SoRa spinning disk confocal with an ORCA Fusion BT sCMOS camera (Hamamatsu) controlled through NIS Elements software (Nikon) (Dema et al. 2024). An Apochromat 40x long working distance (LWD) numerical aperture (NA) 1.15 water immersion objective was used for in-situ imaging of epidermal model (0.16 μm/pixel). For immunofluorescence imaging of epithelial monolayers and mice skin sections, images were acquired on the Yokogawa CSU-W1/SoRa spinning disk confocal system primarily in SoRa mode with a 60 × 1.49 NA oil immersion lens (Nikon), with some datasets collected using an Apochromat 40x long working distance (LWD) numerical aperture (NA) 1.15 water immersion objective. Quantitative analyses were performed using images acquired at consistent magnification within each experiment. 405 nm, 488 nm, 561 nm, and 640 nm solid-state lasers were used at powers between 25-50% (Mayo et al. 2026). Fluorescence images were adjusted for contrast and brightness using ImageJ.

### Image processing and analysis

Image processing and analyses were performed using ImageJ and Imaris.

#### Mouse skin basal layer thickness

Thickness of the KRT14+ basal layer was measured from the KRT14 channel using a line tool using ImageJ at five positions per image within a single z-plane. Measurements from multiple images across 2-3 sections were averaged to obtain one value per mouse.

#### Mouse skin overlap area fraction

Dual-positive KRT14/KRT10 areas were quantified in confocal images with clearly represented basal and suprabasal layers. ROIs were manually defined for each channel in ImageJ, and their intersection was calculated using the AND operator. The resulting overlap area was confirmed to lie within the phalloidin-defined epidermal boundary and normalized to total epidermal area.

#### Mouse skin EdU proportion

EdU-positive nuclei in epidermis in *Fam83h^+/+^* and *Fam83h^-/-^* skin sections were quantified by counting positive nuclei across multiple fields per mouse and averaged per animal. One EdU dataset was excluded due to absence of staining in *Fam83h^-/-^* skin sections, consistent with a technical failure.

#### In vitro keratin immunostaining intensity

The coefficient of variation of keratin signal intensity was calculated in immunostained single cells under SCR and *FAM83H^KO^* conditions using a custom macro in ImageJ. CV was calculated for individual cells and averaged per biological replicate.

#### StrataChip epidermal height quantification

*Epidermal height quantifications*. As described in (Amakor et al. 2026): three random (x, y) coordinates were generated for each device confocal image. The thickness of the epidermis of each point was found and then averaged for each device.

#### StrataChip Thickness of KRT14+ layer

Total epidermal, KRT14, and KRT10 height/thickness was found using the line tool on at least 3 orthogonal projections of the epidermal model. Percent KRT14 thickness of total height was calculated with averaged values.

#### StrataChip Fraction of KRT14/KRT10 volume overlap

Z-stacks were converted to “.ims” files on Imaris. Imaris surface detection tool was used to detect KRT14 and KRT10 positive volume within the image and overlap volume was found using imaris surface to surface statistical comparison.

#### StrataChip p63 positive nuclei

Z-stacks were converted to “.ims” files on Imaris. Imaris spot detection was used to automatically detect nuclei with the DAPI channel and P63 positive nuclei with the P63 channel. Percent P63 was then calculated.

### Statistical analysis

Sample sizes are provided in the corresponding figure legends. Statistical analyses were performed using GraphPad Prism 8. Unless otherwise indicated, graphs display mean ± SD. Differences between two groups were evaluated using unpaired two-tailed student’s t-test and normality was not formally tested. Images are representative of at least three independent experiments unless otherwise mentioned. Experiments were not randomized, and investigators were not blinded during data analysis.

---

**Supplementary Fig. 1.**
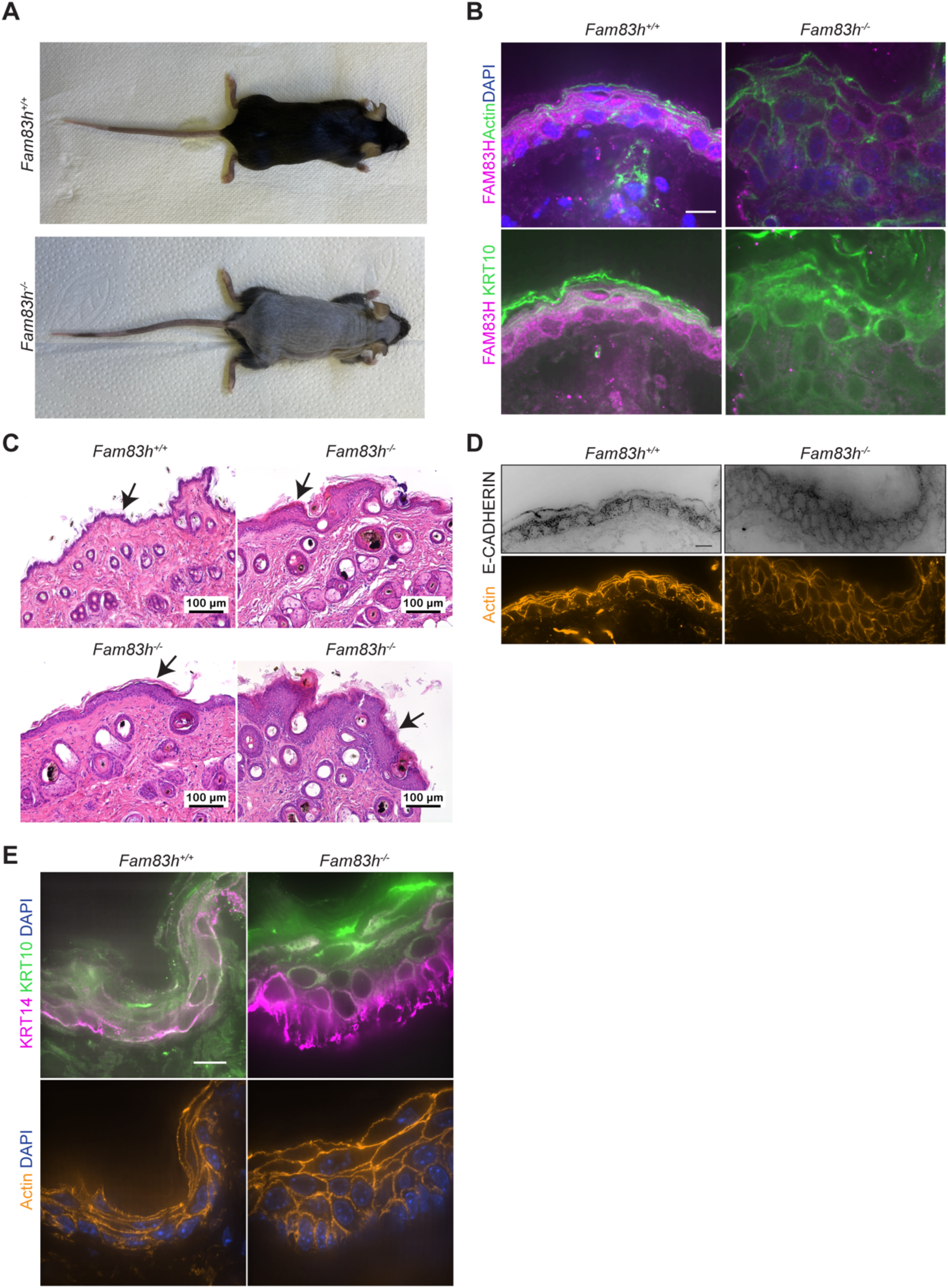
A) Representative dorsal-view images of adult (16 weeks old) *Fam83h^+/+^*and *Fam83h^-/-^* mice. B) Representative z-slice micrographs of back skin sections from *Fam83h^+/+^* and *Fam83h^-/-^* mice at P16, immunostained for FAM83H, phalloidin stained for F-actin (green) and DAPI stained for nuclei (blue) in the top panel and immunostained for FAM83H (magenta) and KRT10 (green) in the bottom panel. Scale bar, 10 μm. C) Representative z-slice micrographs of back skin sections from adult *Fam83h^+/+^*and *Fam83h^-/-^* mice, H&E stained. Black arrows indicate epidermis. Scale, 100 μm. D) Representative z-slice micrographs of back skin sections from *Fam83h^+/+^*and *Fam83h^-/-^* mice at P16, immunostained for E-cadherin (black) in the top panel and phalloidin stained for F-actin (orange) in the bottom panel. Scale bar, 10 μm. E) Representative z-slice micrographs of oral mucosa sections, immunostained for KRT14 (magenta) and KRT10 (green), phalloidin stained for F-actin (orange) and DAPI stained for nuclei (blue) at P16 in *Fam83h^+/+^* and *Fam83h^-/-^* mice. Scale bar, 10 μm.

**Supplementary Fig. 2.**
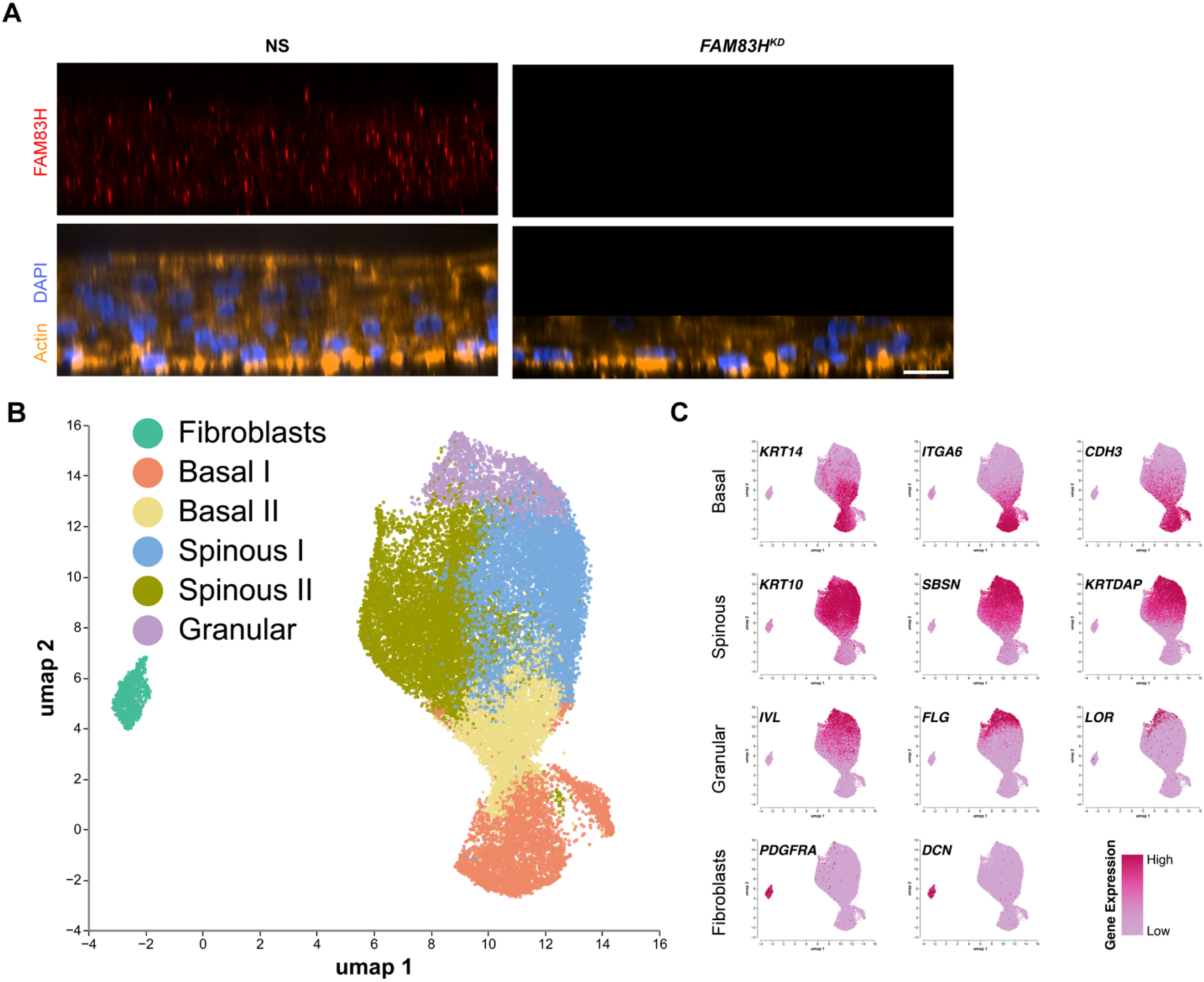
A) Orthogonal slice fluorescence micrograph of 3D epidermal tissue (day 7) immunostained for FAM83H, phalloidin stained for F-actin (orange) and DAPI stained for nuclei (blue). Scale bar, 20 μm. B) UMAP representation of the basal, spinous, granular and fibroblast cell clusters of NS 3D epidermal tissue. C) UMAP with representative gene expression for each layer-specific cell type. Color scale indicates gene expression level from high (dark pink) to low (light pink).

**Supplementary Fig. 3.**
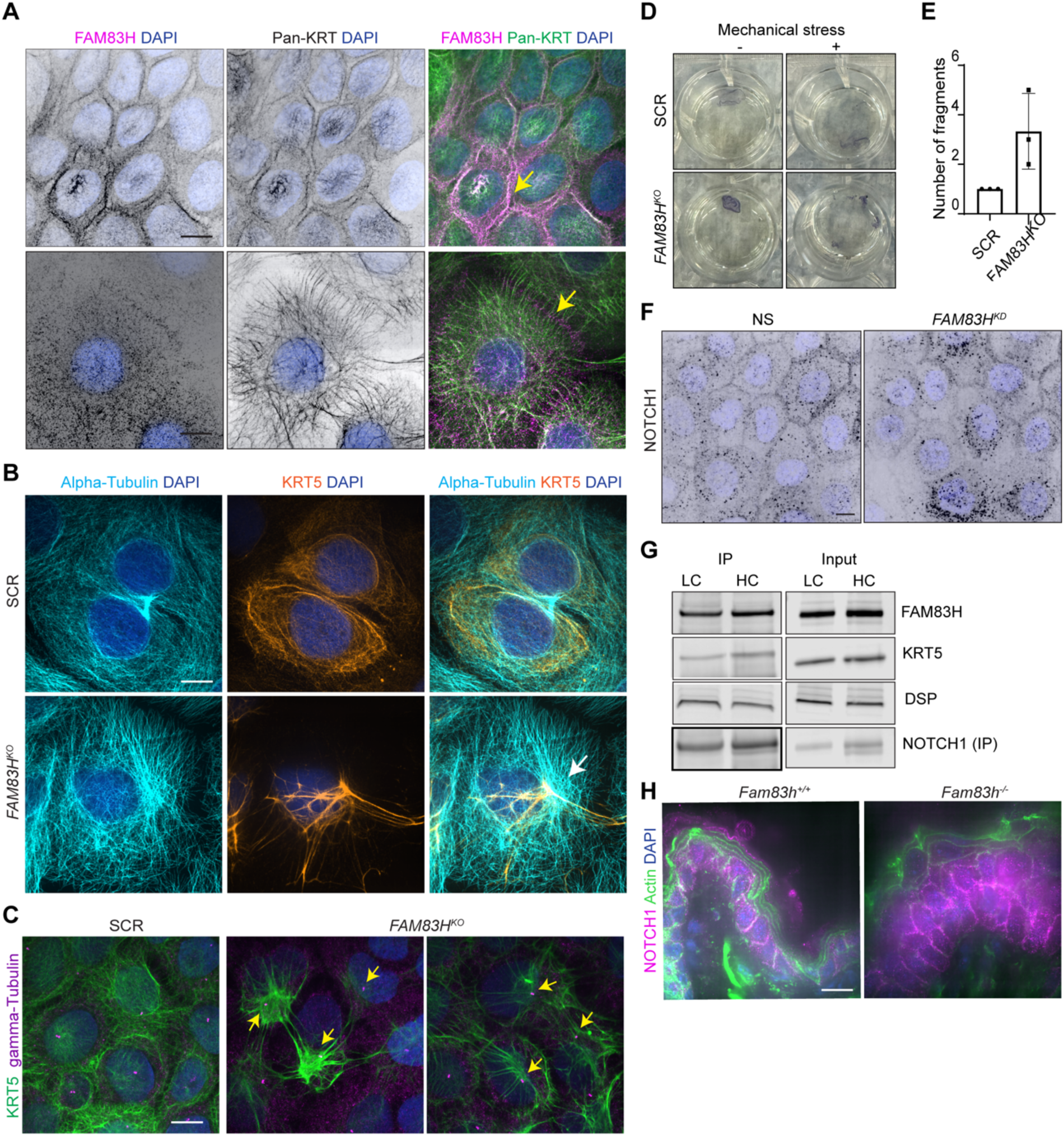
A) Representative z-slice micrographs of HaCaT cells, immunostained for FAM83H and Pan-KRT and DAPI stained for nuclei (blue). Merged images of FAM83H (magenta) and Pan-KRT (green) in the right. Scale bar, 10 μm. B) Representative z-slice micrographs of SCR and *FAM83H^KO^* HaCaT cells, immunostained for KRT5 and alpha-tubulin and merge of Pan-KRT (orange) and alpha-tubulin (cyan). Scale bar, 10 μm. C) Representative z-slice micrographs of SCR and *FAM83H^KO^* HaCaT cells, immunostained for KRT5 and gamma-tubulin and merged images of Pan-KRT (green) and gamma-tubulin (magenta). Yellow arrow indicates the localization of gamma-tubulin. Scale bar, 10 μm. D) Dispase dissociation assay showing the monolayers and fragments of SCR and *FAM83H^KO^* HaCaT cells before and after the shear stress. E) Quantification of number of fragments resulting in SCR and *FAM83H^KO^* HaCaT cell monolayer. N = 3 independent experiments F) Representative z-slice micrographs of cells, immunostained for NOTCH1, DAPI stained for nuclei (blue). Scale bar, 10 μm. G) Western blot of wild-type Ker-CT lysates under low and high calcium conditions followed by NOTCH1 immunoprecipitation, immunoblotted for FAM83H, KRT5, DSP and NOTCH1. H) Representative z-slice micrographs of back skin sections from *Fam83h^+/+^* and *Fam83h^-/-^* mice at P16, immunostained for NOTCH1 (magenta) and DAPI stained for nuclei (blue) and phalloidin stained for F-actin (green). Scale bar, 10 μm.

## Acknowledgments

We thank the members of the Kutys laboratory for insightful discussions. We also acknowledge Joshua Peter (MSKCC) for his assistance with AlphaFold analysis and Olha Fedosieieva for pathological evaluation of H&E slides. This work was supported by grants from the NIH (R35GM150987, S10OD028611). KAJ was supported by Tobacco-Related Disease Research Program Predoctoral Fellowship T33DT6442, and National Science Foundation Graduate Research Fellowship 2038436. BMO, JB and RS were supported by the Czech Academy of Sciences RVO 68378050, and by LM2023036 and CZ.02.01.01/00/23_015/ 0008189 (Upgrade of the large research infrastructure CCP III), co-funded by the European Union and the Ministry of Education, Youth and Sports of the Czech Republic (MEYS). The authors used OpenAI’s ChatGPT and Claude for grammar and language correction during the editing phase of manuscript preparation.

## Notes

### Competing Interest Statement

The authors have declared no competing interest.

### Summary of Updates

Changed one author's order in the file

